# Integrating leaf gas exchange and chlorophyll fluorescence to reveal the long-term regulation of photosynthesis *in situ*

**DOI:** 10.1101/2023.11.22.568237

**Authors:** Jaakko Oivukkamäki, Juho Aalto, Erhard E. Pfündel, Manqing Tian, Chao Zhang, Steffen Grebe, Yann Salmon, Teemu Hölttä, Albert Porcar-Castell

**Affiliations:** Institute for Atmospheric and Earth System Research/ Forest Sciences, Viikki Plant Science Center, Faculty of Agriculture and Forestry, University of Helsinki, Latokartanonkaari 7, FI 00014 Helsinki, Finland; Heinz Walz GmbH, Eichenring 6, D-91090 Effeltrich, Germany

**Keywords:** alternative energy sinks, electron transport rate, field measurements, MICRO-PAM, photorespiration, Silver birch, stomatal conductance

## Abstract

Understanding the diurnal and seasonal regulation of photosynthesis is an essential step in quantifying and modeling the impact of the environment on plant function. Although the dynamics of photosynthesis have been widely investigated in terms of CO_2_ exchange measurements, a more comprehensive view can be obtained when combining gas exchange and chlorophyll fluorescence (ChlF) measurements. However, such integrated measurements have been so far restricted to short term analysis using portable systems that combine IRGA and PAM-ChlF techniques. Here we introduce and demonstrate a new method for integrated, long-term and *in situ* measurements of leaf gas exchange and ChlF, based on an autonomous gas exchange system and a new miniature PAM-fluorometer. The method is used to simultaneously track the dynamics of the light and carbon reactions of photosynthesis at a 20-minute resolution in leaves of silver birch during summer time. The potential of the method is initially demonstrated using the ratio between electron transport and net assimilation (ETR/A_NET_). We successfully captured the diurnal patterns in the ETR/A_NET_ during summer time, including a drastic increase in ETR/A_NET_ upon a high-temperature period. We suggest that these measurements can provide valuable data to model and quantify the regulation of leaf photosynthesis *in situ*.

**Highlight:** We introduce new integrated measurements to help resolve the seasonal and diurnal dynamics of photosynthesis regulation by combining long-term simultaneous measurements of gas exchange and chlorophyll fluorescence in field conditions.

## 1. Introduction

Photosynthesis, and its regulation, have been widely investigated in terms of gas exchange measurements. These measurements are based on enclosing leaves or small shoots inside a cuvette and feeding it with air of known chemical composition. Subsequently, infrared gas analyzers (IRGA) are used to analyze the chemical composition of the air flowing out of the cuvette. Differences in the concentration of CO_2_ between the incoming and outgoing air provide information on the net exchange of CO_2_ (A_NET_) as mediated by the rate of RuBisCO carboxylation (true photosynthesis), RuBisCO oxygenation (photorespiration), and mitochondrial respiration (Farquhar et al., 1980; von Caemmerer and Farquhar, 1981; Lawson et al., 2014). Likewise, changes in the concentration of H_2_O provide information on the rate of transpiration and can be used to estimate stomatal conductance (g_s_) and, by extension, the concentration of CO_2_ in the intercellular air spaces, an essential parameter to model photosynthesis (Von Caemmerer and Farquhar, 1981; Flexas et al., 2002 Flexas et al., 2004). As such, portable IRGA systems have been widely used to measure and model the environmental regulation of photosynthesis, either by following the dynamics of A_NET_ under natural conditions, or by investigating the response of A_NET_ to changes in irradiance (Björkman, 1981; Demmig-Adams and Adams, 1992), temperature (Bernacchi et al., 2001; Sage and Kubien, 2007) or CO_2_ availability (Cornic and Briantais, 1991, Ainsworth and Rogers, 2007). Similarly, long-term automated chamber systems can be deployed in the field to investigate the diurnal and seasonal regulation of photosynthesis *in situ* (Bloom et al., 1980; Hari et al., 1999; Kolari et al., 2014). However, measurements of photosynthetic gas exchange provide only a partial view of the photosynthetic process from the perspective of the carbon reactions, although the regulation of photosynthesis includes also the light reactions (Kramer et al., 2004; Gu et al., 2019).

Chlorophyll fluorescence (ChlF) has been used extensively to investigate the regulation of the light reactions of photosynthesis (Genty et al., 1989; Maxwell and Johnson, 2000; Murchie and Lawson, 2013). Among others, ChlF can be used to estimate the photochemical efficiency of Photosystem II (PSII) and the rate of linear electron transport between PSII and PSI (ETR) which is coupled to the production of ATP and NADPH by the light reactions (Genty et al., 1989; Klughammer and Schreiber, 1994), in addition to multiple other parameters reflecting the energy partitioning between photochemical and non-photochemical processes in the photosystem, as reviewed by Lazár (2015). Together, ChlF and gas exchange measurements can provide a more comprehensive view of the regulation of photosynthesis (Cornic and Briantais, 1991; Laisk and Loreto, 1996; Yin et al., 2011). For example, combined measurements of ChlF and gas exchange have been used to investigate how the production of NADPH is partitioned between alternative pathways and RuBisCO carboxylation/oxygenation reactions (Laisk and Loreto, 1996; Morfopoulos et al., 2014), to study the dynamics of mesophyll conductance (g_m_) (Harley et al., 1992; van der Putten et al., 2018), or to estimate the rate of leaf respiration in the light (Yin et al., 2009). Likewise, variations in the ratio between ChlF-based ETR and net CO_2_ assimilation (ETR/A_NET_) provide information on the internal efficiency of energy conversion between light and carbon reactions of photosynthesis. Recently reviewed by Perera-Castro and Flexas (2023), the ETR/A_NET_ ratio can vary in response to environmental stressors, such as drought (Flexas et al., 2002) or nutrient availability (Bazihizina et al., 2015), mediated by the dynamics of mitochondrial respiration, alternative electron sinks and energy pathways or photorespiration. The ETR/A_NET_ ratio can thus provide an integrated measure of the physiological status of the plant (Perera-Castro and Flexas 2023). Clearly, the combination of ChlF and gas exchange can significantly expand our opportunities to investigate and model the regulation of photosynthesis.

Current portable systems for photosynthesis analysis provide an option for combining gas exchange and ChlF measurements. However, these systems are not suitable for continuous and long-term observation in the field. Likewise, although *in situ* and long-term automated gas exchange measurements (Bloom et al., 1980, Kolari et al., 2014) as well as pulse amplitude modulated (PAM) ChlF systems (Porcar-Castell et al., 2008; Porcar-Castell 2011; Martini et al., 2022) have been available for quite some time, the long-term integration of gas exchange and ChlF measurements has not yet been accomplished. This limitation could now be overcome with the introduction of the MICRO-PAM (Heinz Walz GmbH, Effeltrich, Germany) system, a miniaturized and weatherproof PAM fluorometer for long-term field operation.

In this technical communication, we present a first demonstration of integrated, continuous and long-term PAM-ChlF and gas exchange measurements for application in field conditions. The new MICRO-PAM was coupled to an automated chamber system for combined and long-term measurements of ChlF and photosynthetic gas exchange. The setup was used to follow the regulation of photosynthesis, including ETR/A_NET_, in leaves of silver birch (*Betula pendula* Roth) growing in the top-canopy in a boreal forest during summer 2021. The potential of the measurements is demonstrated by examining and discussing the diurnal patterns of ETR/A_NET_ and its response to high temperatures and water stress during summer.

## 2. Materials and methods

Measurements were conducted in the top canopy of a 60-year-old silver birch located at the SMEAR-II station (Station for Measuring Forest-Ecosystem-Atmosphere Relations) (Hari et al., 2013) in Southern Finland (61°51′ N, 24°17′ E, and 180 m of elevation), between 29 June 2021 and 9 August 2021. This measurement period included two periods of high temperatures, 13-14 July and 26-27 July, hereafter addressed as high-temperature periods. Top canopy foliage was accessed from permanently installed scaffolding towers. Two contiguous and south/south-west facing shoots were selected from the top of the tree and separately used to: 1) install the integrated and autonomous shoot gas exchange chamber and MICRO-PAM fluorometer (Figure 1), and 2) conduct benchmarking measurements using a MONITORING-PAM (hereafter referred to as MONI-PAM) fluorometer (Heinz Walz GmbH). Due to a power failure, we had a 5-day gap between 30 July and 4 August.

**Figure 1.**
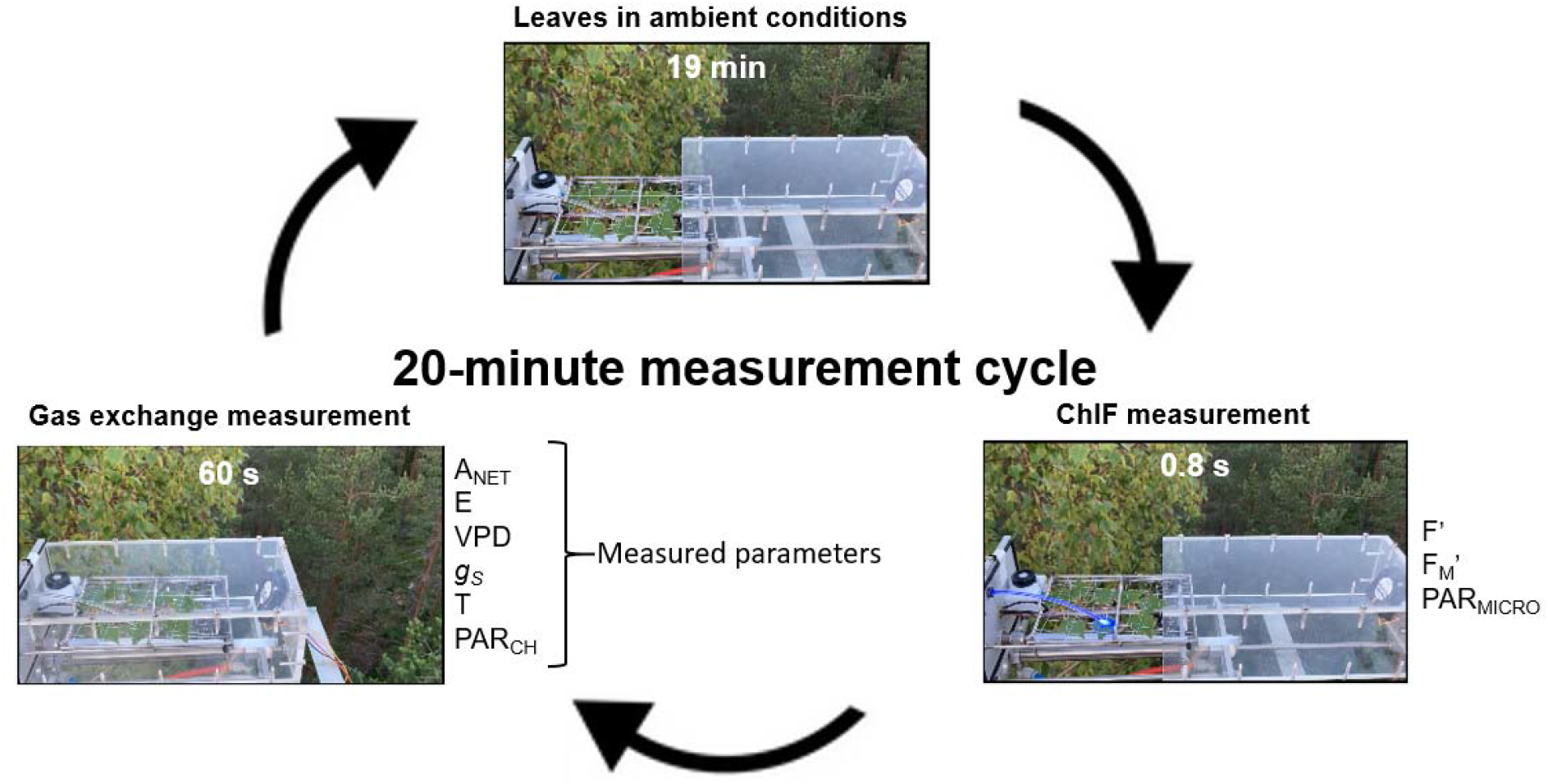
The measurement setup and the 20-minute cycle for the integrated gas exchange and ChlF measurements. Parameters recorded during the cycle are indicated in the diagram and include: current PAM fluorescence yield (F’), maximal fluorescence yield (Fm’), PAR from the MICRO- PAM sensor (PAR_MICRO_), net CO2 assimilation (ANET), transpiration rate (E), vapor pressure deficit (VPD), stomatal conductance (g_S_), as well as chamber temperature (T) and PAR (PAR_CH_).

### 2.1 Long term measurements of shoot-level gas exchange

The shoot chamber used for gas exchange measurements was made of acrylic plastic and had a volume of 2.1 dm^3^. The opening and closing of the chamber was activated with a pneumatically operated system (Figure 1) that operated at a frequency of c. 20 minutes, allowing for regular, automated and long-term measurements of CO_2_ and H_2_O fluxes. Additionally, the chamber housed a PAR sensor (LI-190/R, Li-Cor Inc., Lincoln, Nevada, USA) and a t-type thermocouple for PAR and temperature measurements, respectively. The chamber was closed for 60 seconds every 20 minutes. While closed, gases were sampled via fluorinated ethylene propylene (FEP) tubing using an IRGA system (LI-840A, Li-Cor Inc.) located in a cottage nearby. The chamber enclosed eight leaves arranged on a 2D plane with the help of transparent fishing lines to retain the leaves in a stable position. The full protocol including the ChlF measurements is detailed below. For a more detailed explanation of the shoot chamber system, see Hari et al. (1999) and Kolari et al. (2014).

### 2.2 Long term autonomous measurements of ChlF with the MICRO-PAM and MONI-PAM systems

A MICRO-PAM fluorometer (Heinz Walz GmbH) was installed in the chamber, allowing for simultaneous co-registering of fluorescence parameters. The MICRO-PAM fluorometer is relatively small (13.5 cm x 4 cm x 3.5 cm) and weatherproof, making it compatible with long-term field operation. The system uses blue measuring light and saturating pulses as excitation source. Measuring light and saturating pulses, as well as ChlF are transmitted through a 4.6 cm light guide. In the present study, this light guide was replaced by a longer and curved, 10 cm version to facilitate the measurement of a leaf inside the chamber with the MICRO-PAM attached to the back of the chamber (Figure 1). The tip of the light guide was placed approximately 3-5 mm above the leaf surface at a 35° angle from the leaf plane, measuring an approximate area of 1 cm^2^. The duration of the saturating pulse was set to 0.8 seconds and the intensity to maximum, which delivered approximately 6700 µmol m^-2^ s^-1^ light. The intensity of the measuring light was approximately 1 µmol m^-2^ s^-1^. The detailed ChlF measurement protocol can also be found at dx.doi.org/10.17504/protocols.io.bp2l6xp7klqe/v1. (The DOI link will be activated upon acceptance of the paper, a private link for reviewers can be found in the data availability section. The reviewer link will be removed before publication).

The MICRO-PAM system includes a thermocouple for contact temperature measurements and a quantum light sensor for PAR measurements mounted on a leaf clip. As these variables were already being measured by separate sensors inside the shoot chamber, and since we did not require the leaf clip, the sensors as well as the leaf clip were detached from the MICRO-PAM. The PAR sensor (PAR_MICRO_) was moved to the top of the chamber, back frame and used for quality control during post processing.

Both the MICRO-PAM and MONI-PAM fluorometers were operated through a MONI-DA data acquisition and control unit (Heinz Walz GmbH), housing a micro-SD flash memory card for data storage. The MONI-DA unit can operate multiple PAM fluorometers either using a clock-function or via pre-programmed batch files. In this study, we used the clock function to trigger measurements at a frequency of 20 minutes, to match that of the gas exchange measurements. Originally, the MONI-DA was set to trigger ChlF measurements a few seconds before chamber closure, although the synchronization was lost after a few days (see below). Every 20 minutes, both the MICRO-PAM and MONI-PAM fluorometers registered the current and maximal PAM fluorescence yields, F’ and F_M_’, respectively, allowing for the estimation of the quantum yield of PSII Y(II), the rate of liner electron transport (ETR), and - during nighttime - the maximum quantum yield of PSII or F_V_/F_M_ (see section 2.5).

### 2.3 Reference measurements and ancillary data

The MICRO-PAM time series were validated against those obtained with a MONI-PAM system installed in a nearby shoot. Similar to the MICRO-PAM, the MONI-PAM measures F’ and F_M_’ using a blue measuring light and saturating light pulses (Porcar-Castell et al., 2008). The main differences between the devices are in the geometry of the leaf clip as well as in the fore optics and PAR and temperature measurements. In the MONI-PAM, the measured leaf was attached with a clip ca. 25 mm away from the measuring light, preventing movement of the sample area.

Ancillary data was obtained from the background measurements at SMEAR-II Station. Atmospheric pressure was recorded from a nearby barometer (Druck DPI 260, Baker Hughes, Houston, Texas, USA) at ground level, while precipitation data was gathered using a weather sensor (Vaisala FD12P, Vaisala, Vantaa, Finland) located in a nearby tower at 18 m height. Additionally, soil water content data was collected using soil moisture sensors (Delta-T ML3, Delta-T Devices Ltd, Cambridge, England), averaged from five locations near the measurement tower.

### 2.4 Data analysis and processing

While A_NET_ was directly calculated by the gas exchange system, we calculated saturation vapor pressure (SVP), vapor pressure deficit (VPD) and stomatal conductance (g_s_) from the data gathered with the gas exchange chamber. SVP (kPa) was first calculated using the Magnus-Tetens equation (Tetens, 1930; Monteith and Unsworth, 1994):

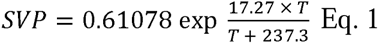

where T is temperature inside the cuvette (°C). Next, we calculated vapor pressure (kPa) using ambient H_2_O concentration (mol/m^3^) and temperature inside the cuvette following the ideal gas law. This allowed us to calculate the vapor pressure deficit (VPD) (kPa), as:

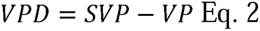

Finally, VPD was used in the calculation of stomatal conductance (g_s_), calculated as:

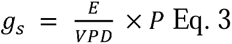

where g_s_ stands for stomatal conductance (mmol m^-2^ s^-1^), E for the transpiration rate (mmol m^-2^ s^-1^) measured by the gas exchange system and P for the atmospheric pressure (kPa).

To calculate the rate of linear electron transport (ETR), we first calculated the quantum yield of PSII (Y(II)), as:

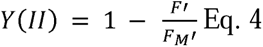

And the subsequently estimated ETR as (Genty et al., 1989):

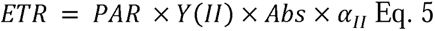

Where Abs is the PAR absorption by the leaf, set here to the default value of 0.84 as a first approximation. The α_II_ is a coefficient that accounts for the fraction of absorbed photons that are associated to PSII antenna, again set to the default value of 0.5, as a first approximation (Maxwell and Johnson 2000), and PAR is the photosynthetically active radiation recorded by the sensor (µmol m^-2^ s^-1^). The maximum levels of nighttime Y(II) were here taken as an estimate of the maximum quantum yield of PSII, or F_V_/F_M_, where F_V_ stands for variable fluorescence (F_V_ = maximal fluorescence (F_M_) – minimum fluorescence (F_0_)). Finally, to provide information on the internal efficiency of energy conversion between light and carbon reactions of photosynthesis, we estimated the ratio ETR/A_NET_.

### 2.5 Data quality control

Despite our efforts to keep the MICRO-PAM and gas exchange measurements in sync, these gradually got out of sync due to a small drift in the internal MONI-DA clock. This drift produced a temporal mismatch between the two measurements of up to 8 minutes. While the effect of this mismatch would be minor if the illumination remains stable during this period, it could lead to discrepancies between the gas exchange and MICRO-PAM data for example under partly cloudy conditions. To identify and exclude measurement pairs (i.e., ChlF and gas exchange data) registered under contrasting PAR levels, we calculated the ratio of PAR measured by the chamber quantum sensor (PAR_CH_) and PAR measured by the MICRO-PAM quantum sensor (PAR_MICRO_), which reflected the illumination conditions at the time of gas exchange and PAM-ChlF measurements, respectively. Although the MICRO-PAM quantum sensor was positioned at the back of the chamber, differences in the ratio PAR_CH_/PAR_MICRO_ can still be used to discriminate measurements collected under diverging light environment. Subsequently, measuring points where the ratio was outside the values of 0.5-2.0 (Supplementary Figure S1) were filtered out and excluded from the analysis.

## 3. Results and Discussion

### 3.1 The environmental response of net CO_2_ assimilation and electron transport rate: diurnal and long-term patterns

Diurnal patterns of variation in chamber PAR and temperature were typical of the boreal summer (Figure 2 A), with very short nights, radiation levels of up to 1700 µmol m^-2^ s^-1^ during daytime and daily temperature fluctuations of 10-15 °C. Diurnal temperature fluctuations were also reflected in the daily pattern of vapor pressure deficit (VPD) and, by extension, in leaf stomatal conductance (g_s_) (Figure 2 B). During sunny days, stomatal conductance started to increase at sunrise in response to PAR, with maximum g_s_ levels before noon, and gradually decreasing towards the afternoon with increasing VPD (but note how this pattern was muted during the second high temperature period in late July, Fig. 2J).

**Figure 2.**
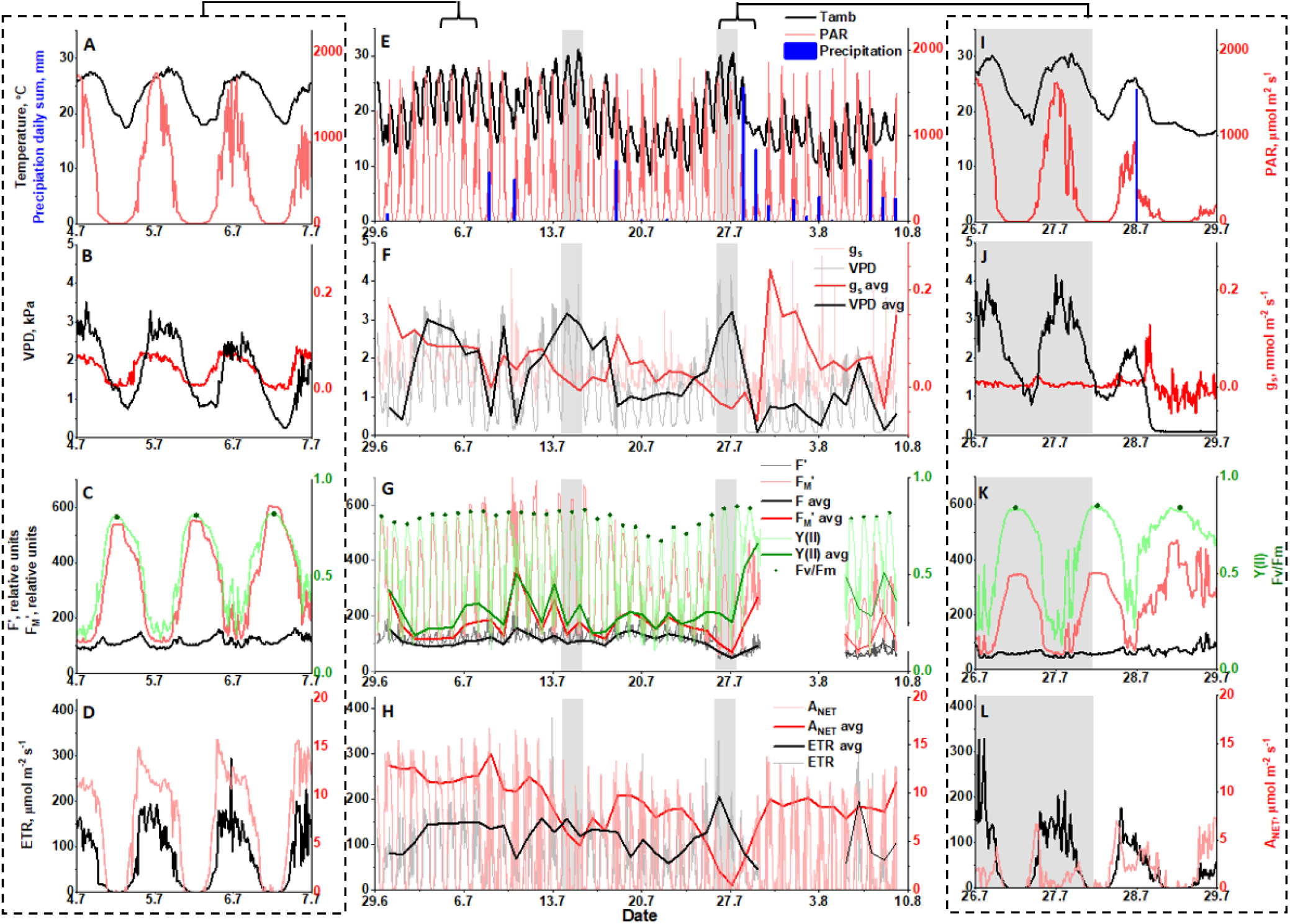
Time series of air temperature, PAR and precipitation (A, E, I); VPD and stomatal conductance (B, F, J); quantum yield of PSII Y(II), maximum quantum yield of PSII or F_V_/F_M_, current (F’) and maximal PAM fluorescence yield (F_M_’) (C, G, K); as well as the rates of linear electron transport (ETR) and net CO_2_ assimilation (A_NET_) (D,H, L), throughout the study period. The left column panels provide a highlight of early July measurements, while the right column panels highlight the second high temperature period. Thick lines in the middle panels denote noon averages (11:00-15:00). Grey shading in middle and right hand columns indicate high temperature periods. The date label presented on the x-axis is situated at noon for each corresponding date.

In turn, the diurnal patterns in the quantum yield of PSII (Y(II)) (Figure 2 C), linear electron transport rate (ETR) and net assimilation (A_NET_) (Figure 2 D), as acquired with our integrated measurements, not only captured the direct influence of PAR and temperature on ETR and A_NET_, but highlighted also the contrasting regulatory dynamics that exist between light and carbon reactions throughout the day. Although both ETR and A_NET_ increased during the morning and decreased towards the evening in response to PAR, the daily patterns of variation were clearly different, with ETR peaking around noontime, whereas maximum A_NET_ was observed in the morning hours, when stomatal conductance was higher (Fig. 2D, C). Daytime A_NET_ values in the beginning of July were around 15 µmol m^-2^ s^-1^, consistent with previous observations for birch in the area (Aalto and Juurola, 2001; Atherton et al., 2017). Finally, the daily variation in Y(II), decreasing around noontime with increasing PAR levels, was consistent with the known activation of regulated thermal dissipation or non-photochemical quenching (NPQ) (Porcar-Castell, 2011).

Throughout the study period, the dynamics in air temperature, PAR and precipitation were indicative of a relative hot summer (Figure 2 E). The mean temperature for July 2021 was 19.0 °C, noticeably higher than the July mean temperature of 16.2 °C for the period 1991-2010 in Hyytiälä (Jokinen et al., 2021). Interestingly, there were two periods when daily temperatures surpassed 30 °C (14-15 July and 26-27 July), which were here used to further emphasize the informative value of our integrated measurements (highlighted with grey in Figure 2). Furthermore, although the amount of precipitation for July was within normal levels, with 97 mm compared to an average of 85 mm for the period 1991-2010 (Jokinen et al., 2021), precipitation was concentrated towards the end of the study period.

Together, the combination of high temperatures (promoting increased evapotranspiration) and limited precipitation resulted in a gradual decrease in soil water contents during the study period (Figure S2), which reached minimum levels during the second high temperature period, just before precipitation partly replenished the soil water storage. These dynamics are consistent with the gradual decrease observed in the daily average *g_s_* from the beginning of the study period until the rainy day on the 28 July (Figure 2E), denoting the stomatal response to the increasingly limited plant water availability and high evaporative demand (Figure 2 F). The regulatory response to decreasing plant water status could be best appreciated during the second high temperature period, where VPD levels of up to 4 kPa combined with minimal soil water availability resulted in *g_s_* values close to zero (Figure 2F,G), which is a typical response to water stress, as the plant tries to conserve water by keeping the stomata relatively closed throughout the day (Martin-StPaul et al., 2017).

Similar to *g_s_*, the daily average A_NET_ gradually decreased during July, and especially during the second high temperature period (Figure 2 L), when net assimilation around noontime was close to zero and clearly limited by stomata. Decreased stomatal conductance would reduce the concentration of CO_2_ inside the leaf, promoting the oxygenation of RuBisCO (photorespiration) relative to the carboxylation (true photosynthesis) (Farquhar and Sharkey, 1982; Flexas et al., 2000). In addition, increasing temperature improves also the affinity of RuBisCO for O_2_, further enhancing photorespiration (Ogren, 1984; Flexas et al., 2002), and increases also the rate of mitochondrial respiration (Busch, 2020), which would have further contributed to decrease in A_NET_ during high temperature periods.

Unlike A_NET_ and g_s_, the diurnal and seasonal patterns in Y(II) and ETR remained relatively stable during the study period and did not present a clear response to the decreasing plant water status. This is consistent with previous observations indicating that PSII activity is not as severely affected by drought as the carbon reactions (Lu and Zhang, 1998; Flexas et al., 1998, 2002; Helm et al., 2020).

A stable seasonal pattern could be also noted in terms of maximal quantum efficiency of photochemistry (F_V_/F_M_), which decreased only in response to the cooler days between the two high temperature periods, but not in response to water limitations. Similar dynamics were recorded by the complementary MONI-PAM dataset (Figure S5). The limited response of F_V_/F_M_ to water stress and its high sensitivity to low temperatures have been described earlier (Oliveira and Peñuelas, 2005; Wang and Huang, 2004). In turn, the decrease of F_V_/F_M_ during the cooler days could be associated with the accumulation of sustained forms of regulatory non-photochemical quenching or NPQ in response to high irradiance and lower temperatures (Porcar-Castell 2011; Verhoeven 2014), which would be consistent with the observed decrease in F_M_’ for the same period (Figure 2G). Overall, the results emphasize the added informative potential of field integrated ChlF and gas exchange measurements whereby we can gain deeper insight into the environmental response of photosynthesis *in situ* by simultaneously following the dynamics of light and carbon reactions.

Noticeably, we observed a sudden decrease in the level of current (F’) and maximal (F_M_’) PAM fluorescence yields recorded by the MICRO-PAM system that took place between two consecutive measuring points during the hot morning of the 27^th^ July at the start of the second high temperature period (Figure 2G and S3A). The decrease did not reverse and levels of F’ and F_M_’ remained lower for the rest of the study period. Such a decrease was however not registered in the supporting MONI-PAM dataset, neither associated with a clear change in the quantum yield of PSII (Figure S3A), which would point out to a technical reason. Since the decrease was not gradual it could suggest that there was a movement of the leaf being measured, decreasing the absolute PAM signal levels. Leaves were kept in place by transparent fishing lines and it is possible that these lines got loose in response to the high temperature, allowing the leaf to move slightly off the fiber field of view. Yet, we did not observe any significant differences in the slope of the relationship between the nighttime F_V_/F_M_ levels from the MICRO-PAM and MONI-PAM systems when comparing the days prior and posterior to this sudden change (Figure S3B), suggesting that the decrease in F’ and F_M_’ had not disrupted the capacity of the MICRO-PAM to estimate the quantum yield of PSII, and by extension ETR. We did however observe a systematic difference in F_V_/F_M_ levels between the two instruments, where MONI-PAM F_V_/F_M_ levels tended to be slightly higher and presenting lower sensitivity to the cooler nights (Figure S5 and S3B). We are unsure on what could be the cause of these differences, but given that both systems used the same blue measuring light, they could be related to the different measurement geometry, imperfect calibration, or simply due to slight differences in the local light environment between leaves.

### 3.2 The potential of *in situ* and integrated measurements of ChlF and gas exchange: the case of the ETR/A_NET_ ratio

The potential of our integrated ChlF and gas exchange measurement setup was further demonstrated by resolving the diurnal and long-term dynamics in the ETR/A_NET_ ratio. The ETR/A_NET_ ratio reflects the internal efficiency of energy conversion between light and carbon reactions of photosynthesis, denoting the amount of electrons that are being transported for every molecule of CO_2_ contributing to the net CO_2_ balance of the leaf (i.e. net photosynthesis). The theoretical minimum value for the ETR/A_NET_ is 4, since 4 electrons are needed for the generation of two NADPH molecules, used in turn for the fixation of a single CO_2_ molecule in the Calvin-Benson-Bassham cycle (Farquhar et al., 1980). Yet, processes such as photorespiration, mitochondrial respiration, or alternative energy sinks will decrease the internal efficiency (Alric and Johnson, 2017; Walker et al., 2020) and values between 8-10 are normal for non-stressed C_3_ plants (Perera-Castro and Flexas, 2023).

To the best of our knowledge the method here reported presents the first *in situ* and long-term time series of the ETR/A_NET_ ratio, obtained here at a high temporal resolution of 20 minutes. During early July (Figure 3, inset), the diurnal variation ranged from 3.2 - 5 at dawn and during sunset, to about 14 - 17 electrons per net CO_2_ assimilated around noon, which is when the ratio peaked. Likewise, the ETR/A_NET_ ratio increased to values of up to 62 and 247, during the first and second high temperature period, respectively, before returning to normal levels (Figure 3). On one hand, these diurnal and seasonal patterns are largely consistent with the contrasting dynamics in Y(II), ETR and A_NET_ discussed above, reflecting the increasing role of photorespiration around noon-time as well as during the high temperature periods, when decreasing stomatal conductance limiting CO_2_ diffusion into the chloroplasts, together with increasing temperature, would shift the partitioning of electrons (via NADPH) into photorespiration (Flexas et al., 2002, D’Ambrosio et al., 2006). On the other hand, the diurnal and seasonal dynamics in the ETR/A_NET_ ratio observed here could also reflect multiple other physiological and methodological factors.

**Figure 3.**
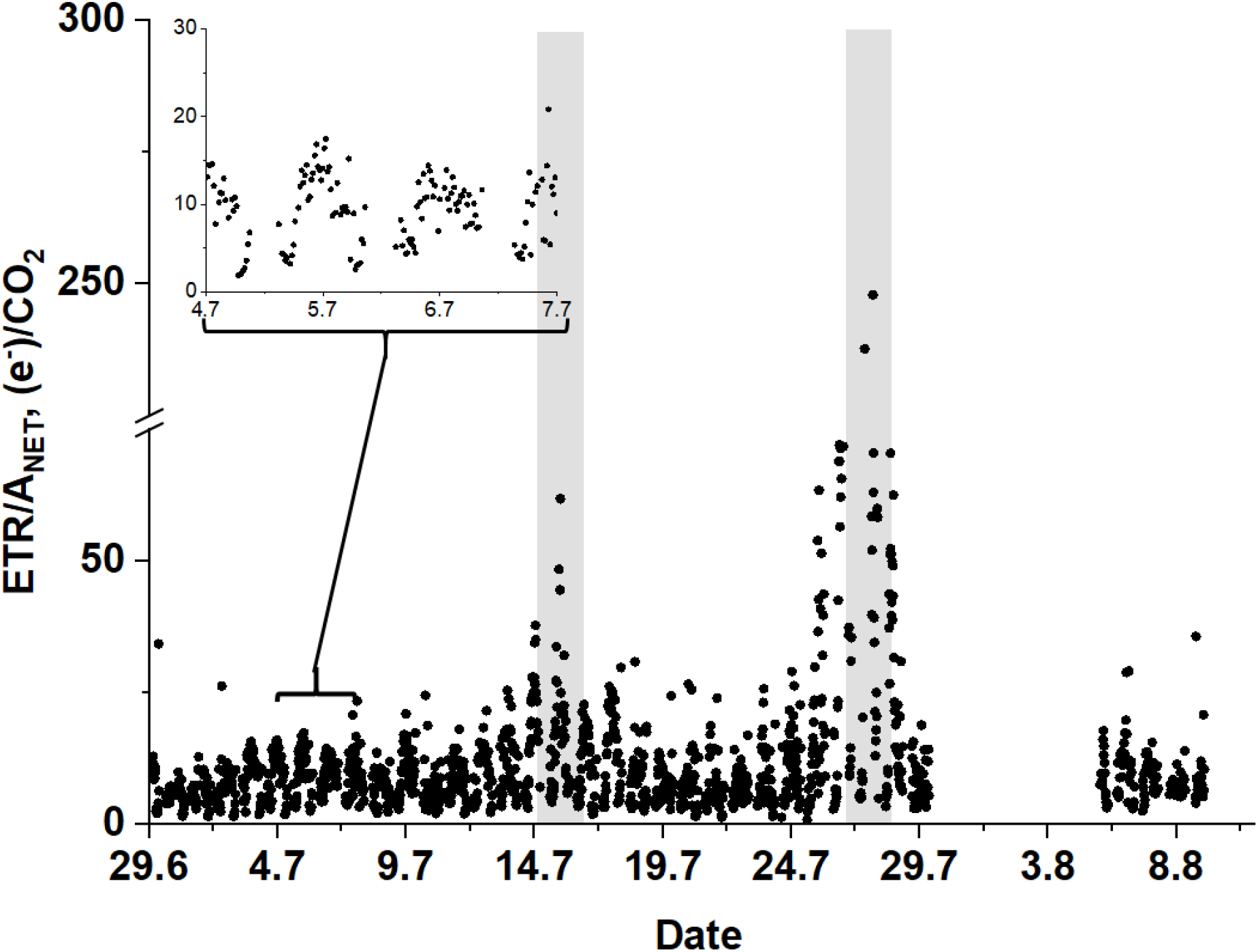
Time series of the ETR/ANET ratio in birch leaves obtained with our integrated ChlF and gas exchange measuring system during the study period. The two high temperature periods are highlighted with grey shading. An example of the diurnal variation can be seen in the inset for illustrative purpose. Note how ETR/ANET values increase around noon and especially in response to the high temperature periods. The date label presented on the x-axis is situated at noon for each corresponding date.

Physiologically, in addition to the dynamics of stomatal conductance reported here, changes in mesophyll conductance (Harley et al 1992; van der Putten et al., 2018), alternative electron sinks (Laisk and Loreto, 1996; Alric and Johnson, 2017), changes in PAR absorption mediated by chlorophyll changes or chloroplast movements (Blache et al., 2011; Pfündel et al., 2018), or even dynamics in the distribution of absorbed excitation energy between PSII and PSI via α_II_, could have all contributed to the observed patterns in ETR/A_NET_. Although a detailed characterization of the sources of variation was outside the scope of the present note, we believe the integrated measurements presented here could prove very useful to further study and model the dynamics of these processes *in situ*, helping elucidate their diurnal and seasonal regulation and their functional significance.

Methodologically, ETR/A_NET_ values during the early morning and late evening hours were clearly below the theoretical minimum, indicating that our measurements were likely underestimating ETR. Possible causes for ETR underestimation could be related to limitations in the cosine response of the PAR sensor, which could have underestimated PAR at low solar elevation, as well as differences in the footprint of ChlF and gas exchange measurements. Our ChlF measurements were here conducted using blue measuring light, which is mostly absorbed by the top layers of chloroplasts in the palisade (Vogelmann and Han, 2000). These chloroplasts are therefore exposed to higher radiation levels and they could be expected to present lower quantum yields compared to chloroplasts located deeper in the spongy mesophyll. In contrast, gas exchange measurements are an integrated measure of the photosynthetic performance of all chloroplasts, which could partly contribute to the underestimation of ETR/A_NET_ levels. Likewise, in the absence of data for our leaves and given the demonstrative purposes of this technical note, we did not correct our F-levels for the contribution of PSI, which can contribute with around 30% to the minimal F_0_ PAM levels in C_3_ plants (Pfündel 1998), and that, if left uncorrected, can result in underestimation to the quantum yield of PSII (Pfündel 1998, Porcar-Castell et al., 2006). Similarly, we used a fixed standard level for PAR absorption (*Abs*) and the fraction of absorbed light allocated to PSII units (α_II_) in Eq. 5, which could also affect the accuracy of our ETR/A_NET_ estimates. Overall, although methods for the estimation of PSI contribution, *Abs* and α_II_ do exist (Genty et al., 1990, Pfündel 1998, Valentini et al., 1995, Yin et al., 2009), methods to continuously track their temporal dynamics in the field await still to be developed.

### 3.3. Final remarks and future steps

The technique presented here offers a relatively straightforward way to upgrade chamber-based field measurements of photosynthesis, either in measuring campaigns or long-term installations, offering a more comprehensive view to investigate the processes underlying the regulation of photosynthesis. For example, the methods here presented can provide very valuable information to investigate and model the temporal dynamics between ChlF and photosynthetic carbon uptake (Gu et al., 2019, Johnson and Berry, 2021), which remains a critical step towards the interpretation of solar-induced fluorescence (SIF) data in terms of photosynthesis (Porcar-Castell et al., 2021, Sun et al., 2023). In the future, the technique could be further improved by: a) replacing the blue PAM measuring light by a red, amber or even green measuring light that penetrates deeper into the leaf (although the signal strength should be checked), to confer a better match between the footprint of the ChlF and gas exchange measurements (Evans et al., 2017), b) improving the synchronization between measurements, which could be attained by controlling the MONI-DA unit from a centralized unit, c) integrating a spectrometer system to provide contiguous measurements of spectral reflectance and fluorescence along with PAM ChlF and gas exchange data, and last but not least, by complementing the measurements with estimations of Abs, α_II_, and the contribution of PSI to total ChlF, in order to provide a more accurate estimate of ETR.

## Supplementary data

The following supplementary data are available at JXB online.

Figure S1 The filter used to remove measurements with differing light conditions.

Figure S2 Changes in volumetric soil water content during study period.

Figure S3 Comparison of ChlF parameters from the MONI- and MICRO-PAM systems.

Figure S4 Daily average temperatures and comparison of F_V_/F_M_ values between the MONI- and MICRO-PAM systems.

## Author contributions

Conceptualization (APC), Investigation (JO, APC, MT, JA), Data analysis and curation (JO, MT, JA, CZ), Writing – original draft preparation (JO, APC), Writing – Review & Editing (EE, SG, YS, TH). All authors contributed to editing and commenting the manuscript towards the preparation of its final version.

## Data availability

All primary data (including metadata) within the manuscript will be openly available on the Zenodo online repository upon acceptance of the manuscript. Note: Private link to the ChlF protocol for reviewers: https://www.protocols.io/private/81BF7235885D11EEBD0B0A58A9FEAC02 to be removed before publication.

